# Modeling stem cell nucleus mechanics using confocal microscopy

**DOI:** 10.1101/2020.08.31.274092

**Authors:** Z Kennedy, J Newberg, M Goelzer, Stefan Judex, CK Fitzpatrick, G Uzer

## Abstract

Nuclear mechanics is emerging as a key component of stem cell function and differentiation. While changes in nuclear structure can be visually imaged with confocal microscopy, mechanical characterization of the nucleus and its sub-cellular components require specialized testing equipment. A computational model permitting cell-specific mechanical information directly from confocal and atomic force microscopy of cell nuclei would be of great value. Here, we developed a computational framework for generating finite element models of isolated cell nuclei from multiple confocal microscopy scans and simple atomic force microscopy (AFM) tests. Confocal imaging stacks of isolated mesenchymal stem cells (MSC) were converted into finite element models and siRNA-mediated LaminA/C depletion isolated chromatin and LaminA/C structures. Using AFM-measured experimental stiffness values, a set of conversion factors were determined for both chromatin and LaminA/C to map the voxel intensity of the original images to the element stiffness, allowing the prediction of nuclear stiffness in an additional set of other nuclei. The developed computational framework will identify the contribution of a multitude of sub-nuclear structures and predict global nuclear stiffness of multiple nuclei based on simple nuclear isolation protocols, confocal images and AFM tests.

## Introduction

All living organisms function in and adapt to mechanically active environments at the levels of the organ, tissue, and cell. Mesenchymal stem cells (MSC) are the tissue resident stem cells of musculoskeletal tissues that, at least in part, regulate the adaptive response to mechanical challenge by proliferating and differentiating into distinct cell types^1^. MSC stem cell differentiation is heavily influenced by the stiffness of the extracellular matrix^2^. For instance, plating MSCs onto soft or stiff substrates can drive MSC differentiation towards adipogenesis or osteogenesis, respectively^3^. The means by which MSC can sense the stiffness of its extracellular matrix comprise an interplay of focal adhesions, the cytoskeleton, and the nucleus^4^. When a MSC is placed onto a stiffer extracellular matrix, the cell will increase its size and the number of focal adhesions to its extracellular matrix^5^, promoting cell traction within the extracellular matrix^4^. As the cell spreads on the extracellular matrix, actin microfilaments tug on the nucleus causing it to stretch and deform^6^. These changes in the nuclear structure are critical for cell function. For example, the nuclear membrane is covered with nuclear pore complexes that are sensitive to cytoskeletal deformations of the nucleus^7^. When these pores are opened, the transcriptional factors such as YAP/TAZ are allowed into the nucleus to regulate gene expression^8^. Further, chromatin itself is responsive to mechanical challenge, as the application of mechanical forces can alter heterochromatin dynamics and organization^9,10^. While signaling events such as YAP/TAZ and DNA changes are areas of active research, probing nuclear mechanical properties in living cells remain challenging.

Quantifying the bulk mechanical properties of the nucleus can be performed via atomic force microscopes, micropipette setups, optical tweezers, or microfluidics^11^. While single-cell level optical methods to measure intra-nuclear deformations are emerging^12^, cellular FE models that can capture nuclear structure and predict nuclear mechanics of many nuclei could provide mechanistic information on cell’s mechanical properties and at the same time, present a time-saving and cost-effective alternative. The stiffness of the nucleus is primarily affected by two nuclear components, LaminA/C and chromatin^13^. LaminA/C is a protein that scaffolds the inner nuclear membrane, adding mechanical stiffness to the nucleus while Lamin B does not contribute to nuclear mechanics^14^. Chromatin is made of compact DNA and histones that occupies the interior of the nucleus and also provides mechanical competence^15,16^. Thus, inclusion of these two components is essential for modeling nuclear mechanics.

Here, we propose and validate a method that uses imaging intensity data from confocal images from LaminA/C and chromatin to determine nuclear mechanical properties. To this end, we developed a computational framework capable of producing confocal-image-based finite element models of an MSC nucleus that replicates the structural configuration of both chromatin and LaminA/C. Finite element models constructed with image intensity based elasticity values were calibrated using experimental AFM measurements of cell nuclei with or without LaminA/C. This model was then tested in its ability to predict the stiffness of two additional test nuclei.

## Data Collection, Modeling, Simulation Setup and Methods

### Cell Culture

MSCs were harvested from the bone marrow of 8-wk old male C57BL/6J mice as previously described.^17,18^ Cells used for the experiments were between passage 7 and passage 11. Cells were sub-cultured at a density of 1,800 cells/cm^2^ and maintained within IMDM (12440053, GIBGO) with 10% FCS (S11950H, Atlanta Biologicals) with 1% Pen/strep (GIBCO).

### Nucleus Isolation

MSCs were scraped off their plates using 9 mL of 1x PBS and centrifuged at 1100 RPM at 4°C with a Beckman Coulter Allegra X-30R Centrifuge. MSCs were suspended within 500 µL hypotonic buffer A (.33M Sucrose, 10mM HEPES, pH 7.4, 1mM MgC12, 0.5% w/v Saponin) and centrifuged twice at 3000 RPM, 4°C for 10 minutes using a Beckman Coulter Microfuge 20R Centrifuge. Cytoplasmic supernatant was aspirated away and the remaining nuclei were resuspended using 100 µL of hypotonic buffer A. Cytoplasmic debris was separated from the nuclei by adding 400 µL of Percoll. The resulting mixture was centrifuged at 10,000 RPM at 4°C for 10 minutes. Nuclei were then plated in a .01 Poly-L-Lysine coated 35 mm cell culture dish and incubated for 25 minutes.

### Nucleus Stiffness Data Using AFM

Force-displacement curves of isolated nuclei were acquired using a Bruker Dimension FastScan Bio AFM. Tipless MLCT-D probes (0.03 N/m spring constant) were functionalized with 10 µm diameter borosilicate glass beads (Thermo Scientific 9010, 10.0 ± 1.0 µm NIST-traceable 9000 Series Glass Particle Standards) prior to AFM experiments using UV-curable Norland Optical Adhesive 61 and, a thermal tune was conducted on each probe immediately prior to use to determine its spring constant and deflection sensitivity. Nuclei were located using the AFM’s optical microscope and engaged with a 2-3nN force setpoint to ensure contact prior to testing. After engaging on a selected nucleus, force curve ramping was performed at a rate of 2 µm/sec over 2 µm total travel (1 µm approach, 1 µm retract). Three replicate force-displacement curves were acquired and saved for each nucleus tested, with at least 3 seconds of rest between conducting each test. Measurements that showed less than 600 nm contact with the nucleus were discarded. Measured force-displacement curves were than exported into Matlab (Mathworks, Natick, MA) to generate a curve of points that reflects the mean of the force to displacement curve as well as the standard deviation of the atomic force microscopy experiments.

### Nucleus Imaging

A singe group of MSC was grown within control conditions and isolated using the methods described above. The chromatin of the nuclei was then stained with Hoechst 33342 while the LaminA/C was stained with mAB 4777 (Abcam). 5 nuclei were then imaged using a Nikon A1 confocal microscope at a rate of .2 µm out of plane and .05 µm in plane resolution.

### Measuring Stiffness of Intact and LaminA/C Depleted Cell Nuclei

As we sought to model nuclear stiffness based on confocal images of LaminA/C and chromatin, we first obtained mechanical properties of cell nuclei isolated from live MSCs with or without LaminA/C. Two groups of MSCs were cultured in growth media (IMDM, 10% fetal bovine serum, 1%Pen Strep). One group received a LaminA/C specific siRNA treatment (siLamin), ceasing LaminA/C mRNA expression in MSCs while the other group was treated with a non-specific control siRNA (siControl). 48h after siRNA treatment, cell nuclei were isolated, plated onto 0.1% Poly-L-Lysine coated plates for adherence and subjected to AFM testing to obtain force-displacement curves as we reported previously (**Fig. 1a**)^19^. As shown in immunolabeled nuclei images (**Fig. 1b**), isolated control nuclei appeared round and maintained intact LaminA/C (red) and DNA (blue) confirmation. Shown in **Fig.1c**, force-displacement curves for siControl (N=30) and siLamin (N=73) groups were obtained by shows that the maximum force measured at the AFM tip for the siLamin group was, on average, 59% smaller than the siControl group (p<0.05, Fig. 1c), confirming that nuclei are softer without LaminA/C^14^.

**Figure 1.**
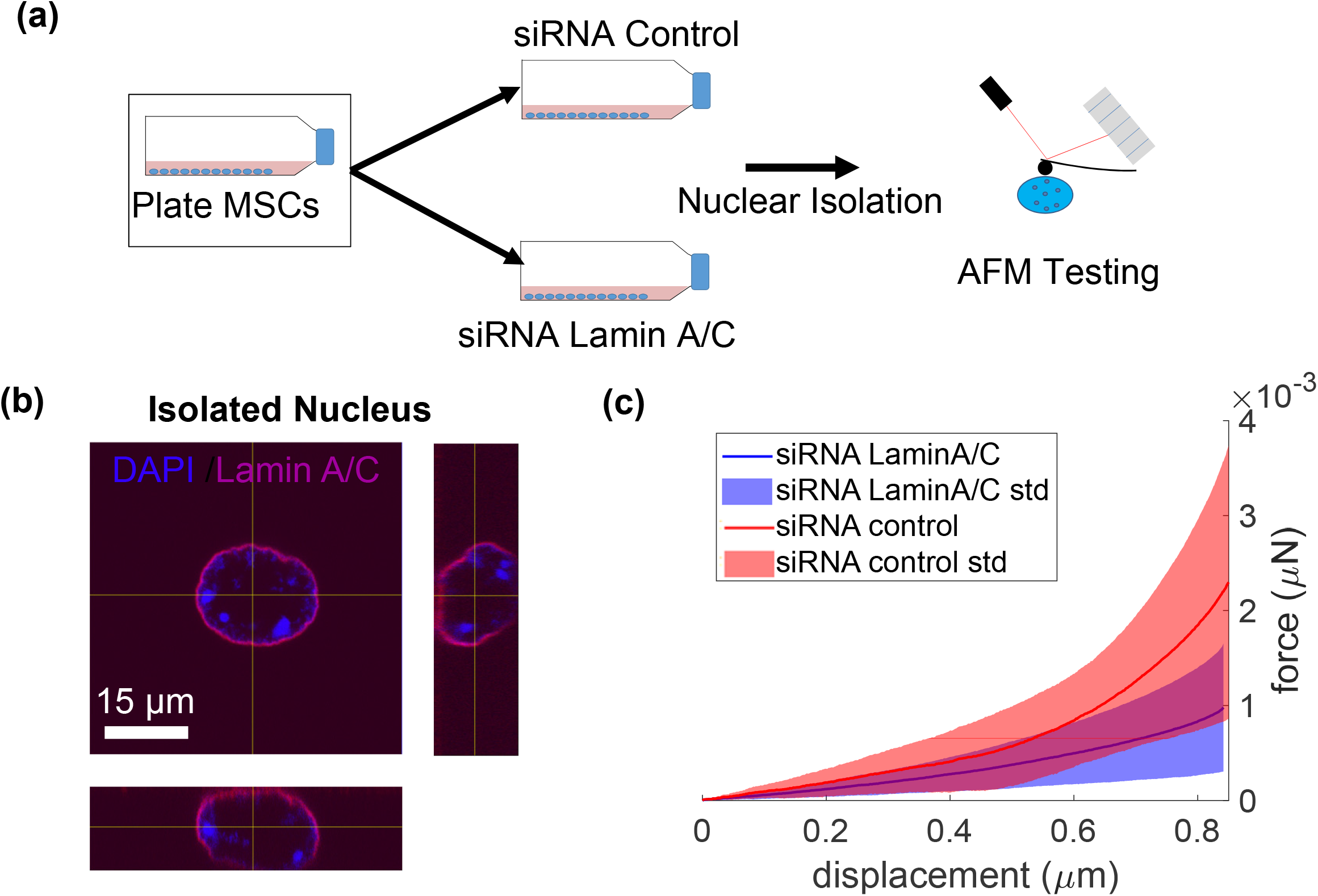
siRNA mediated depletion of LaminA/C decreases isolated nuclei stiffness. a) Two groups of MSCs were grown in 10 % fetal bovine serum. One group received the LaminA/C specific siRNA treatment while the other group was treated with control siRNA. Nuclei were isolated and subsequently subjected to AFM testing. Nuclei of both the control group (n=30) and the laminA/C knockdown group (n=73) were indented by 1 µm using a spherical tip with a diameter of 6 µm. b) Confocal microscopy images of a nucleus stained for chromatin (Hoechst 33342) and laminA/C (cell signaling mAB4777). c) Average (± SD) force-displacement curves for control nuclei (red) and laminA/C siRNA nuclei (blue). The average force-displacement curve for each group is shown as a solid line; less than one standard deviation is shown as shaded area. The purple area represents the overlap of the red and blue areas.

### Mesh Generation from Confocal Scans

To model the contribution of LaminA/C and chromatin separately, we generated two volumetric meshes from each nucleus image. The first mesh was generated using the chromatin signal of the nucleus image and the second one was generated using the LaminA/C signal of the nucleus image. For chromatin, the 3D confocal image of the chromatin was imported into Amira software (ThermoFisher, MA) and the nucleus geometry was manually segmented (**Fig. 2a**). A surface mesh was created that employed triangular S3 elements surrounding the nucleus geometry (**Fig. 2b**). This surface mesh was then imported into Hypermesh (Altair Engineering, MI) to create a volume mesh with C3D4 tetrahedral elements (**Fig. 2c**). The resulting volume mesh was imported into Bonemat software (http://www.bonemat.org/) (**Fig. 2d)** to overlay the volumetric mesh onto the original confocal image and to assign stiffness values to each tetrahedral element using the average voxel intensity (HU) within each element (Equation 1):

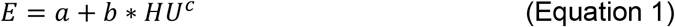

**Figure 2.**
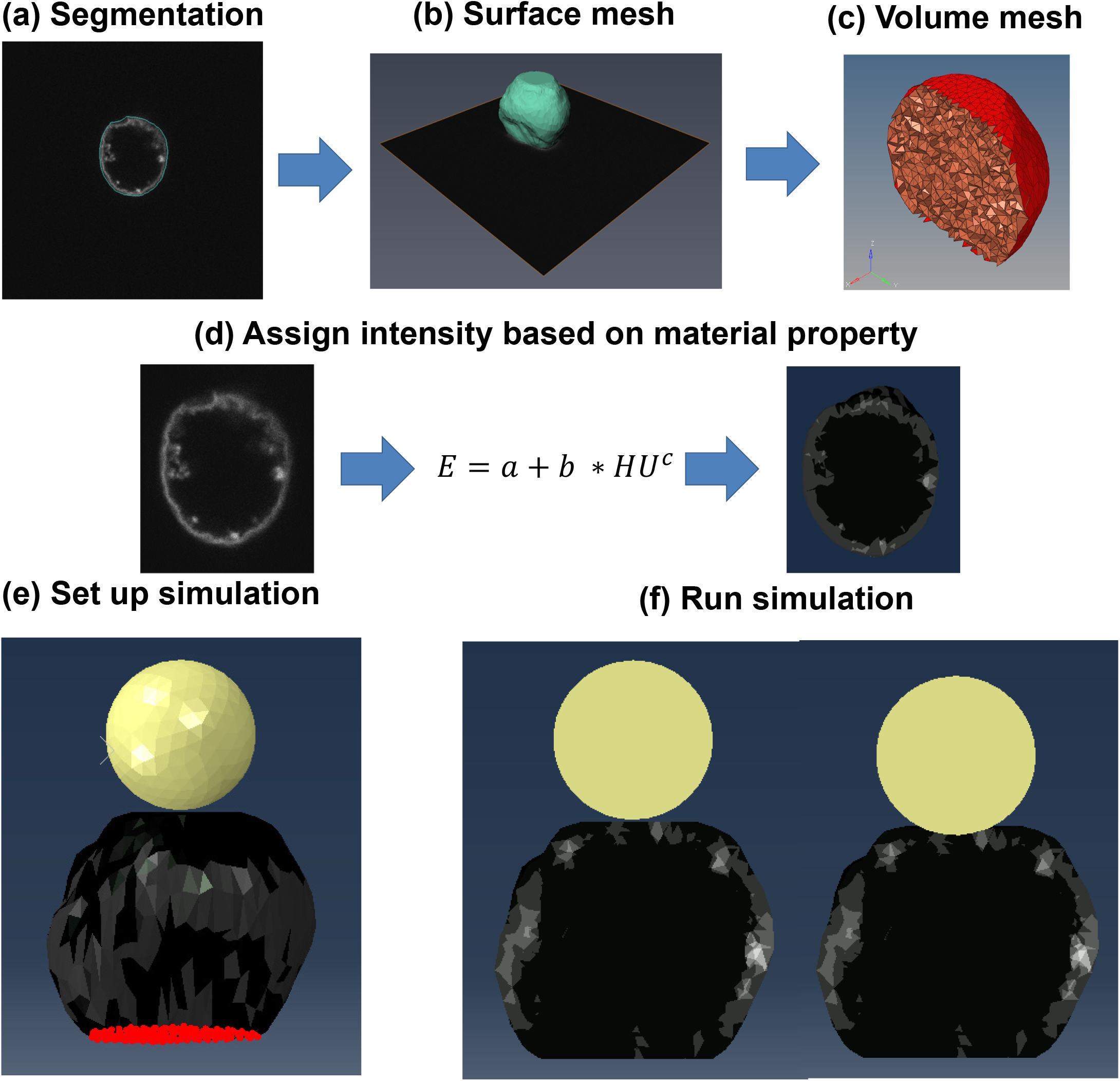
Generation of image-based nucleus model. a) Images of MSC nuclei were manually segmented using Amira to isolate the nuclear geometry. b) Segmented images were then used to create a surface mesh of the nucleus geometry. c) The surface image was converted into a volumetric mesh. d) The volumetric mesh was assigned material properties using the voxel intensity of the original image and the shown equation. e) Image of simulated atomic force microscopy (AFM) experiment with the AFM tip shown in yellow, the heterogeneous nucleus in blue, and the encastered base nodes in red. f) Cross sectional images of the nucleus model before compression (left) and after compression (right).

The term ***a***, representing the intensity-independent elastic modulus, was set to “0” to eliminate any contribution to elasticity outside the image intensity. Terms ***b*** and ***c*** are a set of conversion factors defined during each experiment. Here, we used a linear isotropic elastic material definition with a Poisson’s ratio of 0.5 for each model^20^. For this study, FE meshes were generated for five isolated nuclei. Two representations of each nucleus were generated; one that included both LaminA/C and chromatin, and one which included only chromatin. All the nuclei were imaged via a Nikon A1 confocal microscope with an image depth of .2 µm and a voxel width of .05 µm.

To generate a model that contains both LaminA/C and chromatin (i.e. LaminA/C + chromatin mesh), two identical nucleus geometries were produced. Chromatin meshes were assigned elasticity values using the siLamin nuclei force-displacement curves (i.e. no LaminA/C present). While siRNA procedure does not deplete the entire LaminA/C protein levels, a large decrease in measured force value (**Fig.1c**) shows a substantial decline. The effectiveness of this siRNA procedure was confirmed in a recent publication^21^. Conversion factors for the LaminA/C meshes were derived via utilizing the chromatin mesh elasticity values and the the AFM data for intact nuclei (i.e. siControl nuclei with both Lamin A/C and chromatin present). The chromatin and LaminA/C elasticities in each element were then linearly added to produce the siControl model containing the elasticity of both structures.

### Replicating AFM Experiments *in silico*

Atomic force microscopy simulations were performed in ABAQUS software (2019, Dassault Systemes, France). A replica of the AFM test setup was modeled *in silico* (**Fig. 2e**). The bottom node layer of the nucleus model (red) was fixed to a rigid plane in all orthogonal directions to simulate the nucleus being attached to the poly-L-Lysine coated plate surface. A simulated AFM tip (yellow) was created by positioning a sphere (r=5 µm) made of C3D4 elements with a rigid body material definition above the nucleus model. Contact between the nucleus and the AFM tip was defined as a no-friction contact pair. During simulation, the AFM tip was lowered onto the nucleus until 1 µm indentation was reached (**Fig. 2f)**. The required force to indent the nucleus was recorded. The recorded force-displacement curves were used to quantify the root mean squared error (RMSE) between experimental and *in silico* conditions.

### Determination of the Element Volume for Nucleus Models

To determine the dependence of the AFM indentation force on the volume of the mesh elements, nucleus models were constructed from 5 chromatin nuclei images with element volumes of 5, 4, 3, 2, 1, .8, and .6 µm^3^. The models were assigned temporary elasticity values using the original chromatin images with conversion factors of *a* = 0, *b* = 20, and *c* = 1. The term *a* was set to 0 because of the assumption that there is no base elasticity independent of image intensity, *b* was set to 20 and *c* was set to 1 to scale elasticity *linearly* to image voxel intensity. A representative image for the meshes of nucleus#1, with varying element volumes and with the original images at each mid-orthogonal plane is depicted in **Fig. 3a**. Each nucleus model was subjected to *in silico* AFM experiments. For each nucleus model, the force generated at 1 µm of indentation was recorded and plotted against element volume for each nucleus **(Fig.3b)**. The mean maximum force value and standard deviation started to plateau for element volumes of 1 µm^3^ or smaller, indicating this volume that can be used without affecting the maximum force output (green line).

**Figure 3.**
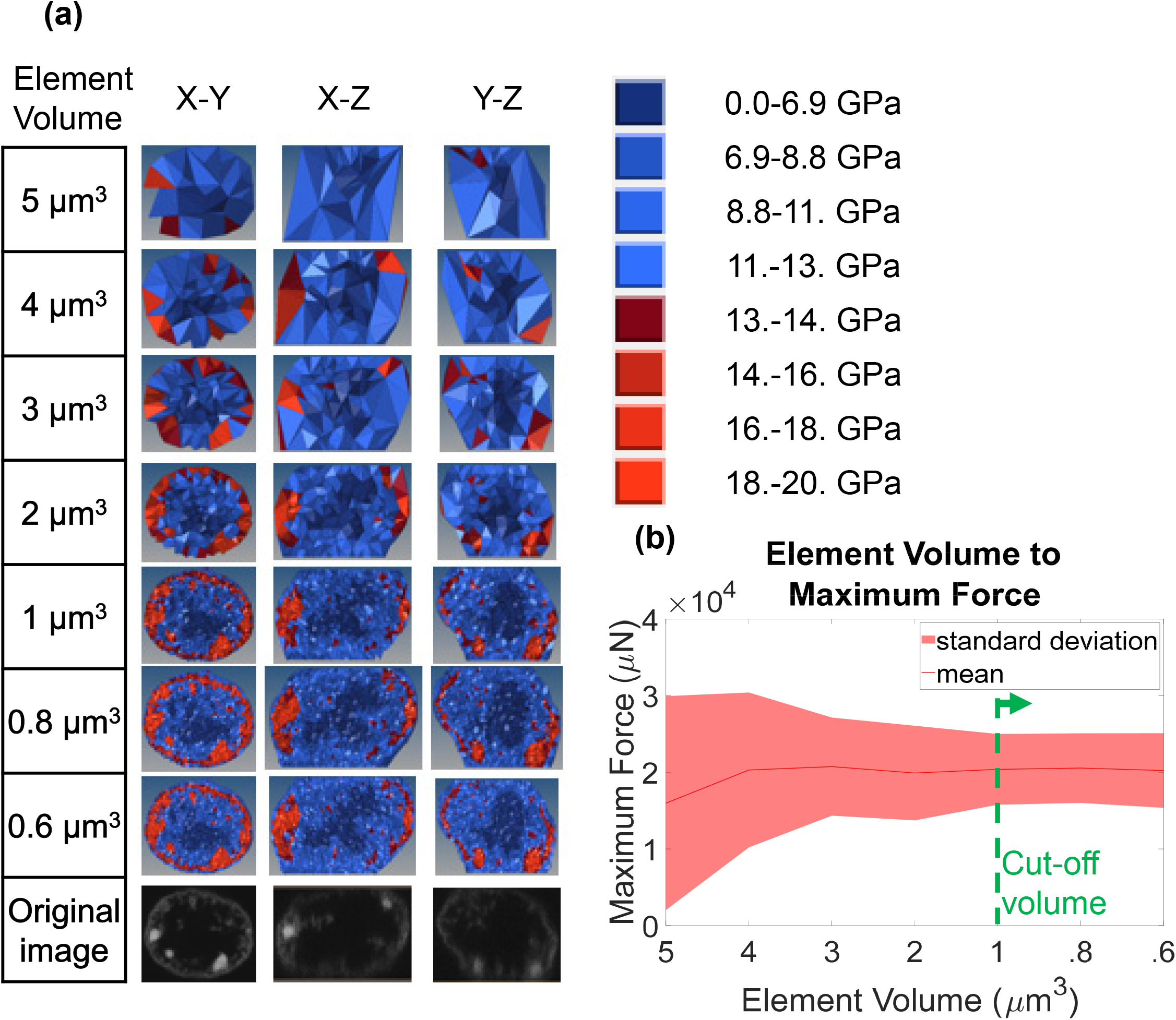
Element size sensitivity analysis. a) Cross sectional images of nuclei models created with elements that have an average element size of of 5, 4, 3, 2, 1, .8, and .6 µm^3^. Material parameters were set to b=20 kPa and c=1. Color maps indicate the corresponding stiffness values. b) Graph of how maximum force, measured at the AFM tip pressing onto the nucleus, versus the element size averaged for three nuclei. The solid line represents the mean and the shaded area indicates the area within one standard deviation. Element sizes smaller than 1 µm^3^ did not affect maximum force and standard deviation (green dashed line).

### Sensitivity of Image Noise to Element Volume

While force sensitivity analysis revealed a cut-off at 1 µm^3^, we also sought to quantify how well element volumes represented the spatial information from confocal images, as this may be important for discerning nuclear deformation patterns. To accomplish this, we only used a chromatin mesh without LaminA/C. Chromatin images for a single nucleus image (Nucleus #1) was converted into six finite element models meshed with average element sizes of 3, 2, 1.5, 1, .8, .6, and .3 µm^3^ and assigned elasticity values using the temporary conversion factors *a* = 0, *b* = 20, and *c* = 1. A Matlab script extracted a 3D image from each mesh with the 2D image from a transverse plane (Z=7 µm) (**Fig. 4**, top row).

**Figure 4.**
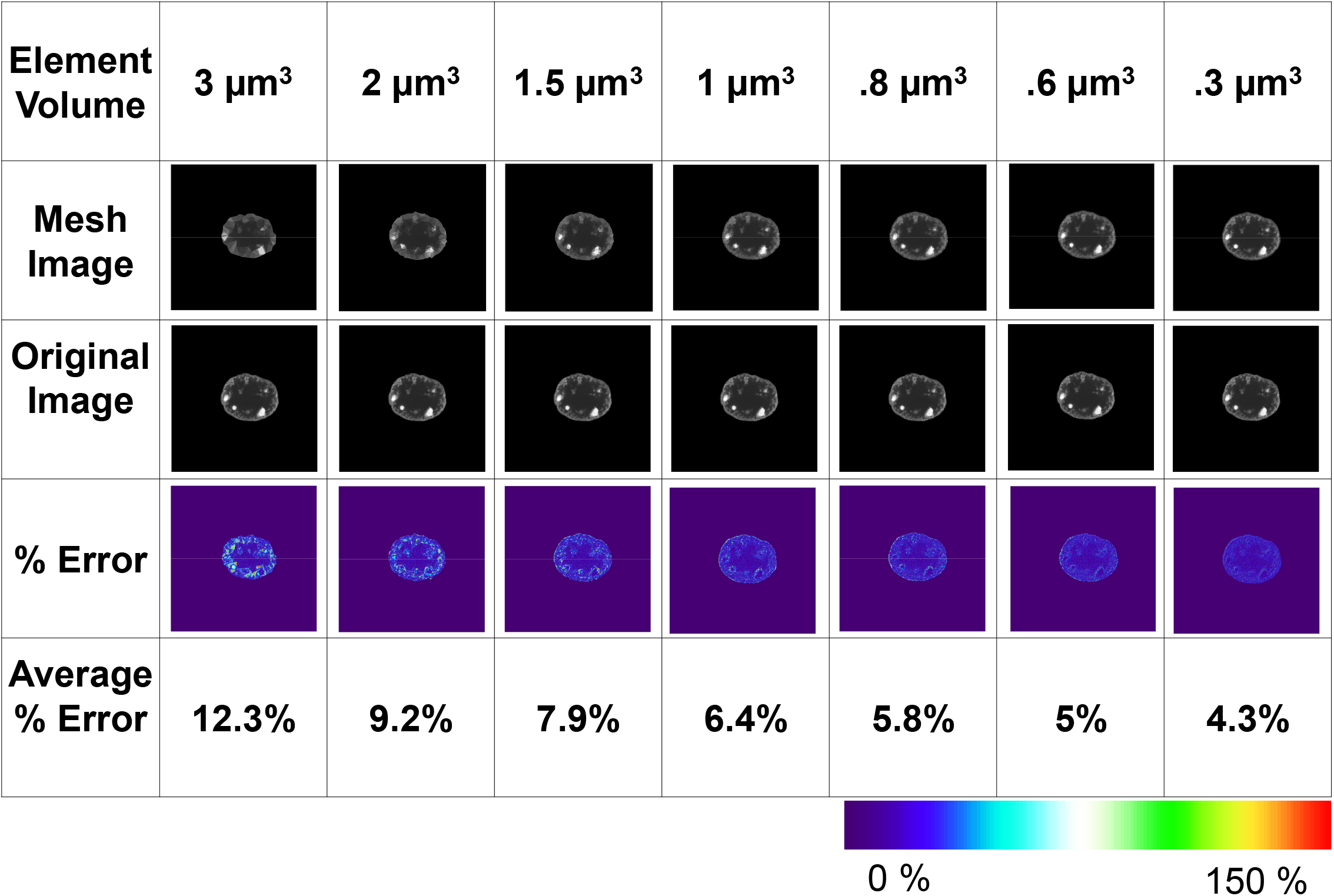
Element size error analysis. Representative sagittal plane images with element volumes of 3, 2, 1.5, 0.8, 0.6, and 0.3 µm^3^ (2^nd^ row) were compared against the matching location in the original confocal image (3^rd^ row). Quantification of the pixel-by-pixel intensity values were represented by a % change heat map (4^th^ row). Average % error in 3 and 0.3 µm^3^ element volumes were 12.3% and 4.3%, respectively.

These images were then superimposed onto the original image (**Fig. 4**, second row) and the intensity of each voxel was compared to each voxel in the original image, producing a color map indicating the percent difference (**Fig. 4**, third row). Microscopy noise in the confocal images was accounted for by comparing the average intensity of the DNA free region of interest to each voxel within that region (**Fig. S2**). This analysis produced an average error value of 13%, indicating the amount of inherent noise in the confocal images. This value was subtracted from each voxel to quantify the non-noise related error. These corrected voxel errors were then averaged to generate a final error value (**Fig.4**, bottom row). At 3 µm^3^ element volume, the average error was 12.3%. As element size decreased, the % error also decreased. At 1 µm^3^, the average error was 6.4%. For 1 µm^3^ down to 0.3 µm^3^, the average error only changed by 1.9% indicating a similar cut-off range where 1 µm^3^ voxel volumes can represent 93.6% of the chromatin configuration.

### Response Surface Generation

To identify the sensitivity of the RMSE between experimental and *in silico* force-displacement curves to different combinations of the b and c terms, two response surfaces were generated; first, a coarse 8 × 8 matrix that was followed up by a finer 10 × 10 matrix. The c values were spaced linearly while the *b* values were spaced logarithmically because of the much greater change of b value, that minimized error, compared to the c value. To generate the RMSE between experimental and *in silico* force-displacement curves, all five nuclei confocal microscopy scans were converted to finite element models with an average element size of 1 µm^3^. Each model was assigned elasticity values using the original image and the conversion factors for the given datapoint. All five nucleus models then underwent a simulated atomic force microscopy experiment where force-displacement data from the first 1 µm of nucleus indentation was collected. The resulting force displacement curves were compared to the mean atomic force microscopy curve taken from the experimental AFM indentations, calculating the RMSE. The RMSE of all 5 models were averaged to create each point of the response surface.

### Conversion Factor Optimization

Nucleus models #1, #2, and #3 were selected and converted to finite element models with an element volume of 1 µm^3^. The matlab algorithm “fmincon” was set to use an “SQP” optimization algorithm with constraint/step tolerance set to 1×10^−9^ µN/µm^2^. The *c* term was then constrained to either *c* = 1 for linear material conversion or *c* = 0 for homogeneous material value. A value of *b* = 1E-10 µN/µm^2^ was used as a starting point. This algorithm optimized the *b* value until the optimization constraint/step tolerance was met.

### Statistical Analysis

Results were presented as mean ± one standard deviation (SD) unless indicated in figure legends. For comparisons between groups, a non-parametric two-tailed Mann-Whitney U-test was used. P-values of less than 0.05 were considered significant.

## Results

### Linear relationship between voxel intensity and material property is sufficient for assigning material properties for both SiLamin and SiControl models

To determine the best set of conversion factors for creating nuclei models containing a chromatin mesh only, an 8 × 8 response surface was generated to compare the simulated AFM results to experimental AFM data for the LaminA/C depleted nuclei (**Fig. 5a)**. The *b* values were logarithmically spaced between 1×10^−9^ µN/µm^2^ and 1×10^−3^ µN/µm^2^ and *c* values were linearly spaced between 0.5 and 5. The error associated with each *b-c* combination was found by generating the root mean squared error between the simulated and experimental AFM data. Results showed that for every *c* value, there was a *b* value that minimized the error. To expand on this finding, we selected the two *c* values of 0.5 and 1.1 that produced a minimum value within our original 8 × 8 grid (green dotted boxes). Shown in **Fig.5b**, plotting a refined 10 × 10 response surface around these two *b-c* values, a minimum error along a straight line for different *b* values was visible (dotted red lines), suggesting that minimizing the error was independent of the initial *c* value. Shown in bottom right, setting *c*=1 produced a similar set of *b* values that minimized the error between the simulated and the real AFM experiments, indicating that a linear conversion between pixel intensity and modulus of elasticity could be used.

**Figure 5.**
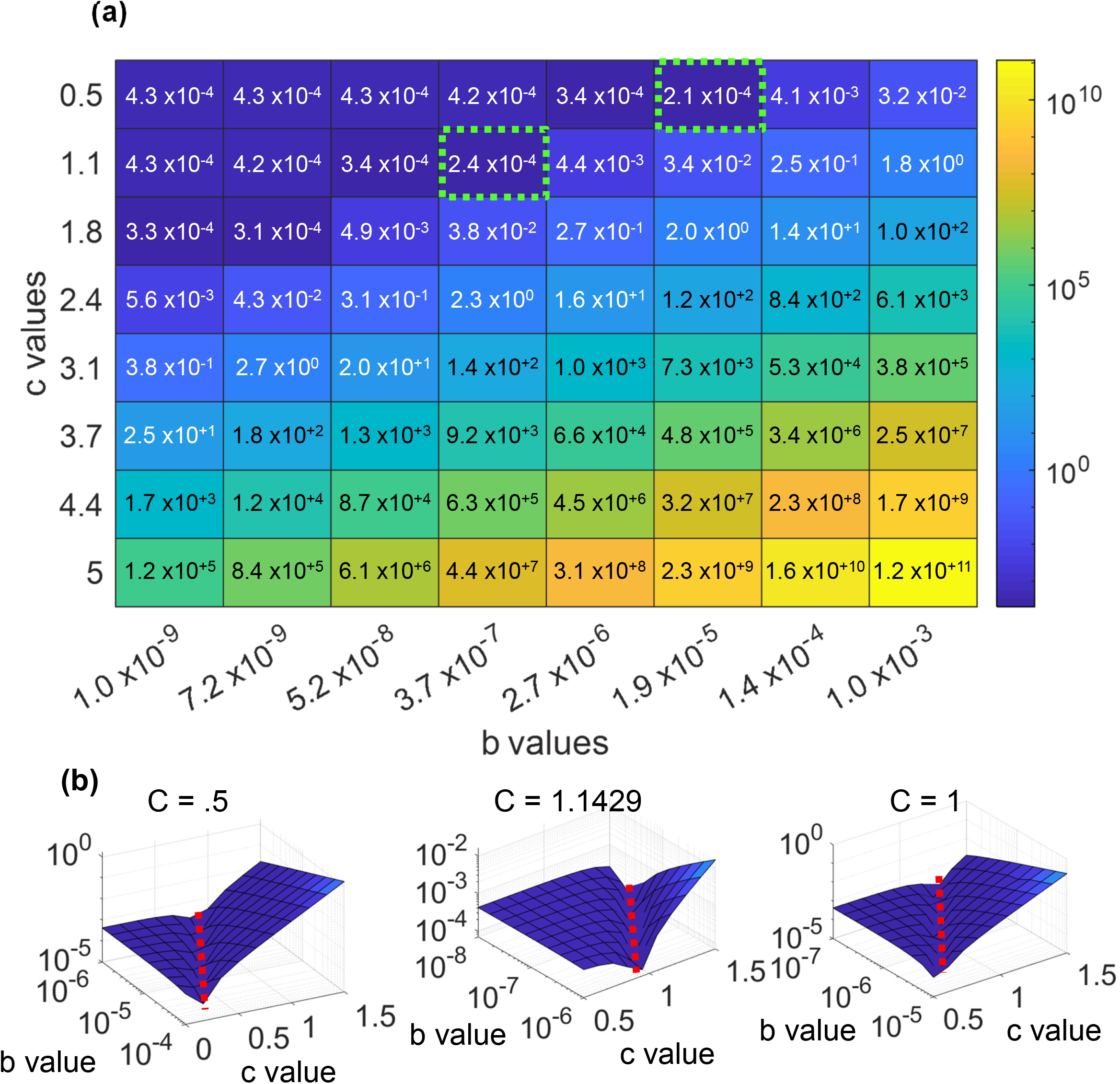
Optimization of LaminA/C knockdown nuclei shows a linear elasticity relationship. a) Error surfaces for 3 LaminA/C depleted nuclei showed a rut-like error when using different b and c values. b) Higher resolution error surfaces were generated around the lowest points of the original surface. These error surfaces produced minimum values on the order of 10^−4^, similar to the error surface generated around c=1, demonstrating that there is a correlation between b and c.

**Figure 6.**
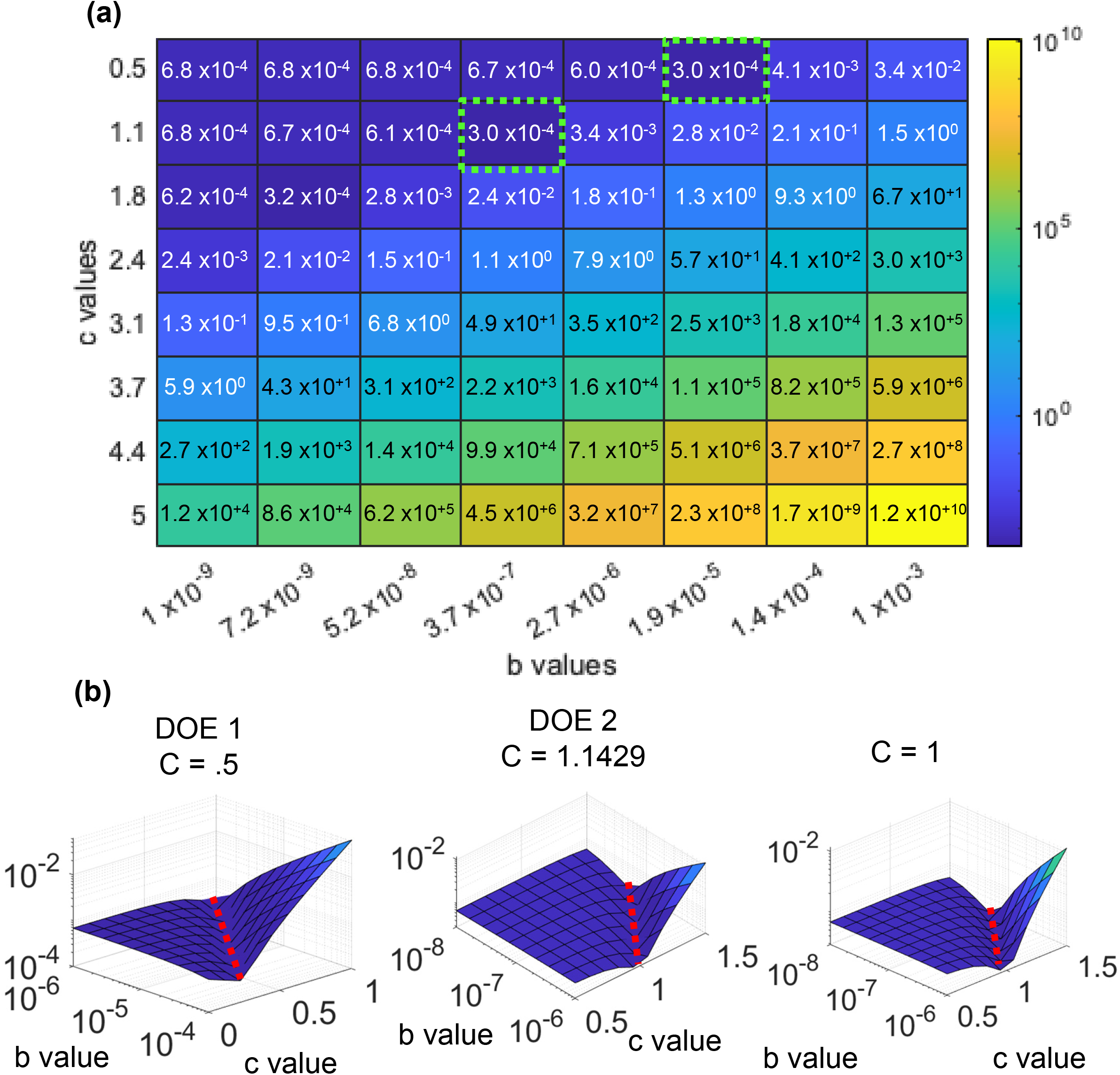
Optimization data for control nuclei knockdown using linear and exponential conversion factors. a) Error surfaces for 3 control nuclei showed a rut like error when using different b and c values. b) Higher resolution error surfaces were generated around the lowest points of the original surface. These error surfaces produced minimum values on the order of 10^−4^, similar to the error surface generated around c=1, demonstrating that there is a correlation between b and c.

Repeating the same procedure using the LaminA/C + chromatin mesh and AFM data from intact nuclei exhibited a similar outcome. We again found that the two *c* values of 0.5 and 1.1 produced a minimum value within our original 8 × 8 grid (green dotted boxes). For the first minimum value, a 10 × 10 surface centered on *b* = 3.7 x10^−7^ µN/µm^2^ and *c* = 1.1 was generated. For the second minimum, we created a surface centered on *b* = 1.9 x10^−5^ µN/µm^2^ and *c* = 0.5. Both surfaces showed a minimum error along a straight line for different *b* values (dotted red lines). Comparing these values to another 10 × 10 surface centered on *b* = 1 x10^−7^ µN/µm^2^ and *c* = 1 showed a similar pattern, indicating that a linear relationship between voxel intensity and material property is sufficient for LaminA/C nuclei. We then set *c*=1 and used the Matlab algorithm “fmincon” optimization algorithm with a step tolerance of 1 x10^−9^ to find the *b* values that minimized the root mean square error for three “training” nuclei (nuclei 1, 2 and 3) for both chromatin and LaminA/C groups. This step resulted in an optimized *b* value of 6.3 x10^−7^ µN/µm^2^ with an error of 5.5 x10^−5^ µN/µm^2^ for chromatin. For LaminA/C, the b mean value was 8.64 x10^−7^ µN/µm^2^ with an error of 3.1 x10^−4^ µN/µm^2^.

### Linear conversion model is distinct from a homogeneous model for chromatin

To test the differences between the homogeneous and linear-elastic heterogeneous models, the homogeneous chromatin models were made from the chromatin structures of nuclei #1 - #3 by setting all the elements to same elastic modulus. The modulus value was determined via minimizing the RMSE between the load-displacement curves of the *in silico* and experimental AFM data of the LaminA/C depleted nuclei, producing a modulus of elasticity of 2.7 x10^−4^ µN/µm^2^ with a RMSE value of 6.2 x10^−5^ µN/µm^2^. There was no statistical difference between the error of the homogeneous and linear-elastic heterogenous models (p=.83). Similarly, applying the error-minimized *b* values to homogenous and heterogeneous models generated from the test nuclei (#4 and #5) resulted in RMSE values of 6.2 x10^−5^ µN/µm^2^ and 5.5 x10^−5^ µN/µm^2^ with similar error values (p=.63), suggesting that the bulk nuclei response can be modeled using either homogenous or heterogeneous models (**Table 1**).

**Table. 1:** Chromatin Material Optimization Data

|  | Linear conversion | Homogeneous model |
| --- | --- | --- |
| B factor/elasticity | $6.3 \times 10^{-7} \mu\text{N}/\mu\text{m}^2$ | $2.8 \times 10^{-4} \mu\text{N}/\mu\text{m}^2$ |
| Average error of testing set | $5.5 \times 10^{-5} \mu\text{N}/\mu\text{m}^2$ | $6.2 \times 10^{-5} \mu\text{N}/\mu\text{m}^2$ |

Next, the *in silico* cross-sectional Von Mises stress during the 1 µm indentation of the tip was compared between the homogeneous and heterogeneous chromatin models of nuclei #4 and #5. To compare the average Von Mises stresses between heterogeneous and homogeneous model simulations, average Von Mises stresses at mid-sagittal planes were plotted and compared across a 1 µm region of interest located between nuclear heights Z=5 µm and Z= 6 µm. The von mises stresses from each element within the models of each group were plotted and statistically compared between the two groups.

The heterogenous models of nuclei #4 (top) and #5 (bottom) showed higher peaks at the nuclear periphery of the region of interest **(Fig. 7b)**. Quantification of the peripheral peak stresses showed 16% greater stresses in the heterogenous model when compared to the homogenous model (p<0.001). The heterogeneous model also showed more efficient load carrying as shown by lower peak stresses that were distributed among more elements when compared to the homogenous model where a smaller number of elements had to carry greater loads (**Fig. 7c)**.

**Figure 7.**
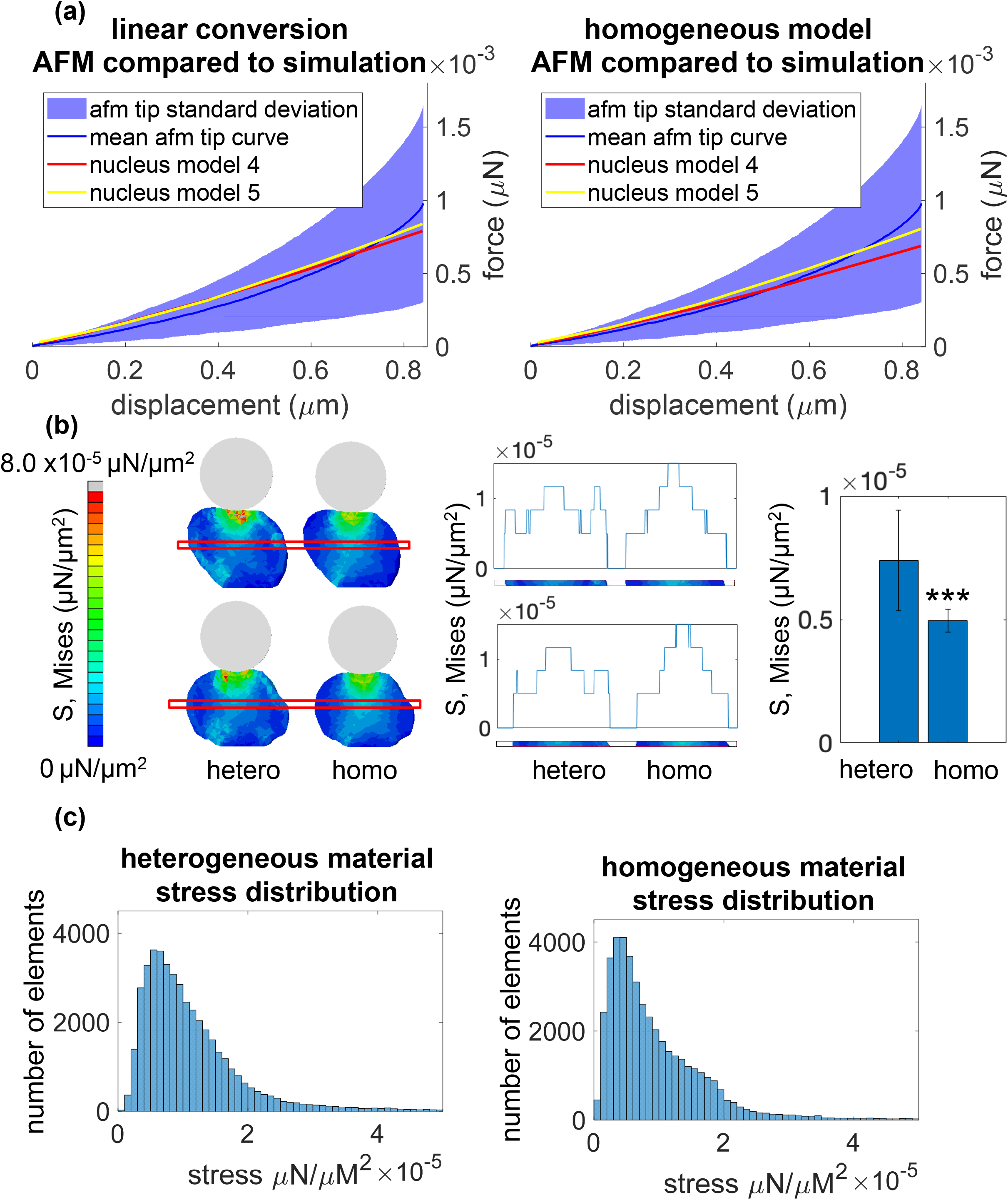
Linear conversion vs homogeneous model for chromatin. a) The simulated force curves were superimposed onto the LaminA/C KO results, comparing the resulting force curves from the linear conversion (left) to the results of the homogeneous model (right). b) Cross-sections of the model when fully compressed were imaged (left) and the average stresses within a 1 µm tall region beginning at a height of Z=5 µm were plotted (middle). Stresses within the outer 25 percentile of both nuclei were plotted with a bar plot (right), showing the difference between the stress distributions within the homogeneous and heterogeneous models. c) Von Mises stress data was collected from the elements of both models at maximum AFM tip compression and plotted within a histogram to show the difference between the stresses developed within the homogeneous and heterogeneous models.

## Discussion

Deformation of the nucleus regulates gene transcription via altering both DNA confirmation^22^ and the nuclear entry of transcription factors such as YAP/TAZ^23^. Nuclear deformation in response to mechanical forces is modulated by the mechanical stiffness provided by chromatin and LaminA/C within the nucleus^13^. The computational framework developed here was able to capture geometrical and structural inhomogeneities of both LaminA/C and chromatin from confocal images. Using constants derived from the calibration of AFM voxel-intensities to elastic moduli, mechanical behavior of nuclei were predicted merely from images without performing a physical mechanical test. The inherent limitation of this approach is that prior to predicting nuclear mechanical properties, a relatively large sample of AFM and confocal images is necessary. Further, while it was outside of the scope of the current study, errors associated with experiment-to-experiment variability of confocal images will need to be evaluated in future studies. Finally, for these predictions to be accurate, the nucleus has to be isolated from the cell as the cytoskeletal contribution to mechanical properties obtained from AFM cannot be avoided in intact cells. Even with these limitations, our method enables the prediction of nuclear stiffness and intra-nuclear deformation with only a simple nuclear isolation protocol and confocal imaging. The mechanical models of isolated, standalone nuclei developed here may also provide mechanistic insight into cellular mechanics and provide a basis for developing mechanical models of nuclei in intact cells in the future.

Our model provides a number of advantages over finite element analyses of the cell nucleus that tend to model the nucleus as a homogenous material with idealized geometry.^24,25^ While comparisons between homogenous and heterogenous nuclear structures showed no significant changes in the “bulk” structural response under in silico AFM experiments, stresses throughout the nuclear structures were different where stresses concentrations were dependent upon the chromatin and LaminA/C distribution density obtained from the original images. As chromatin condensation has been shown to change due to external nuclear loading^26^, these models may provide useful predictions of which regions of chromatin are experiencing larger loads.

Another potential advantage of this modeling system is the incorporation of nuclear envelope proteins into the generated models. In this study, we modeled LaminA/C as a heterogeneous material. Interestingly, the levels of LaminA/C within the nucleus have been shown to change under microgravity^23^. With our model it may be possible to predict the changes in nuclear stiffness due to alterations in LaminA/C levels. Further, the structural contributions of other nuclear envelope proteins such as nuclear pore complexes can also be incorporated into these models in the future, providing a robust computational framework for studying the forces on a number of nuclear proteins.

Previous research described the nucleus’s mechanical elasticity as either linear elastic or hyperelastic^20^. During our experiments we chose to model the nucleus as linear elastic. As both homogenous and linear conversion models of nucleus #4 and #5 produced linear force-displacement curves, we also implemented hyperelastic Mooney-Rivlin and Neo-Hookean material definitions^20^ which again produced linear force-displacement relationships (Fig. S2), suggesting that the shape of *in silico* loading curves were independent of the use of hyperelastic Mooney-Rivlin and Neo-Hookean models. Corroborating these in silico findings, as shown in Fig. S5, 38% of the AFM-tested nuclei showed linear loading curves.

In summary, we generated individual finite element models of nuclei from confocal images. Importantly, these models were tuned to match experimental AFM results, generating a similar bulk mechanical behavior when compared to a homogeneous nuclear structure. We also demonstrated that if a proper relation between chromatin stiffness and image intensity is found, our method can be used to model internal chromatin dynamics within the nucleus. Ultimately, our study may lead to more effective techniques and insight into mechanobiological phenomena within the cell nucleus, elucidating cell nucleus plasticity in response to the application of mechanical forces.

## Authors’ contributions

**Kennedy Z** data analysis/interpretation, manuscript writing, final approval of manuscript.

**Newberg J** data analysis/interpretation, data analysis.

**Goelzer M** data analysis/interpretation, data analysis, final approval of manuscript.

**Judex S** data analysis/interpretation, manuscript writing, final approval of manuscript

**Fitzpatrick CK** financial support, data analysis/interpretation, final approval of manuscript

**Uzer G** concept/design, financial support, data analysis/interpretation, manuscript writing, final approval of manuscript

## Ethics approval

All methods were carried out in accordance with relevant guidelines and regulations of Boise Institutional Animal Care and Use Committee and Institutional Biosafety Committee. All procedures were approved by Boise State University Institutional Animal Care and Use Committee, and Institutional Biosafety Committee.

## Consent for publication

All authors consent to publication

## Availability of data and material

The datasets generated and/or analyzed during the current study are available from the corresponding author on reasonable request.

## Code availability

The code generated and/or analyzed during the current study are available from the corresponding author on reasonable request.

## Acknowledgements

This study was supported by NASA ISGC NNX15AI04H, NIH R01AG059923, and 5P2CHD086843-03, P20GM109095, P20GM103408 and NSF 1929188. We also greatly appreciate the AFM expertise from Dr. Paul Davis and the Surface Sciences Laboratory.

## Supplementary Information

### Funding support

NASA ISGC NNX15AI04H, NIH R01AG059923, and 5P2CHD086843-03, P20GM109095, P20GM103408 and NSF 1929188.

**Figure S1.**
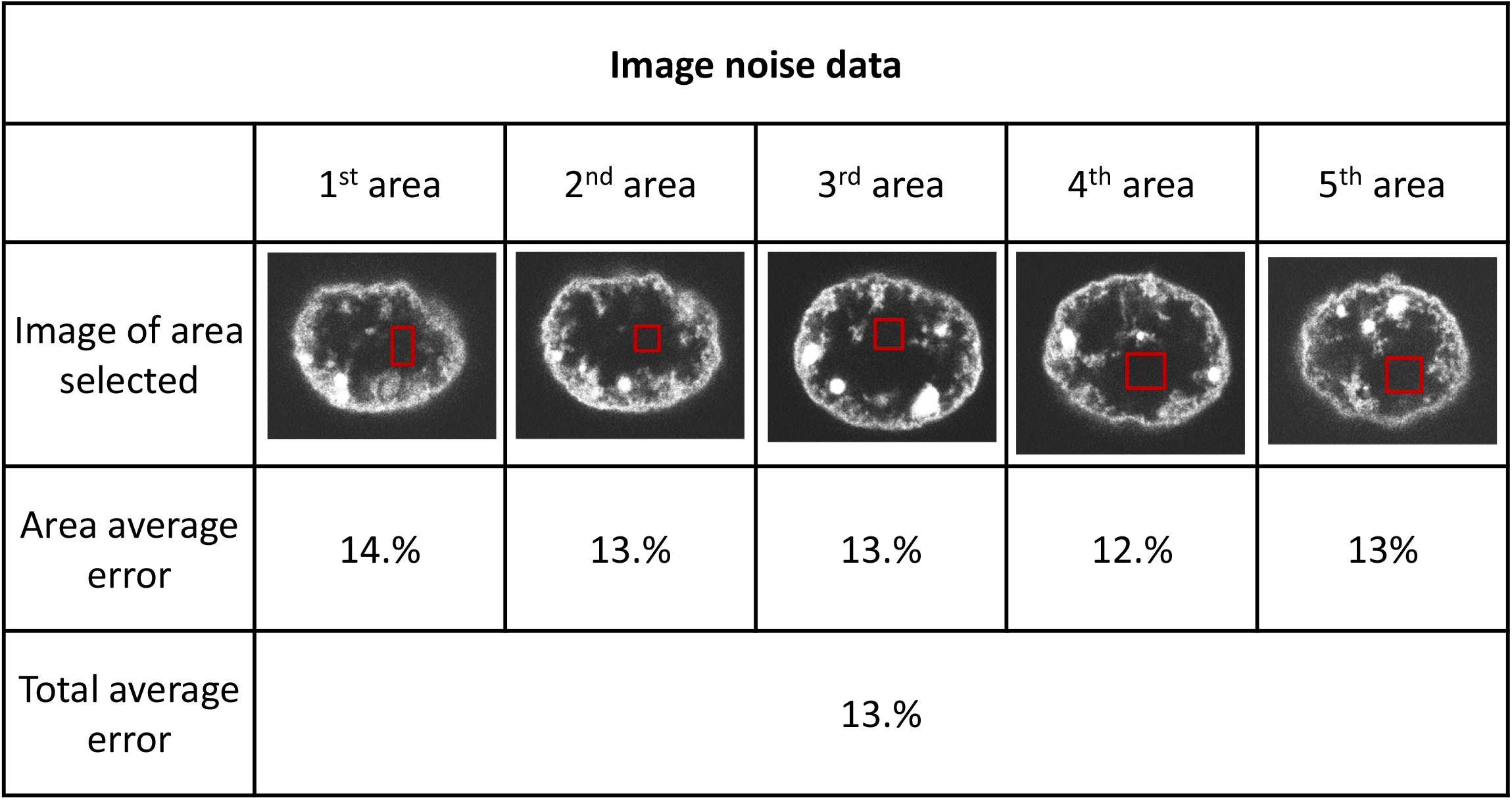
Image Noise. 5 confocal images of chromatin from nucleus 1 were separated and a homogeneous section of the image was then selected from each image. This section was used to quantify the image noise within the microscope by finding the average voxel intensity within the images and comparing this value to the voxels within each area. The error between the average voxel intensity and the accompanying area voxels was averaged to produce an average error of each area. This data was averaged to produce the average noise within the chromatin image of nucleus 1.

**Figure S2.**
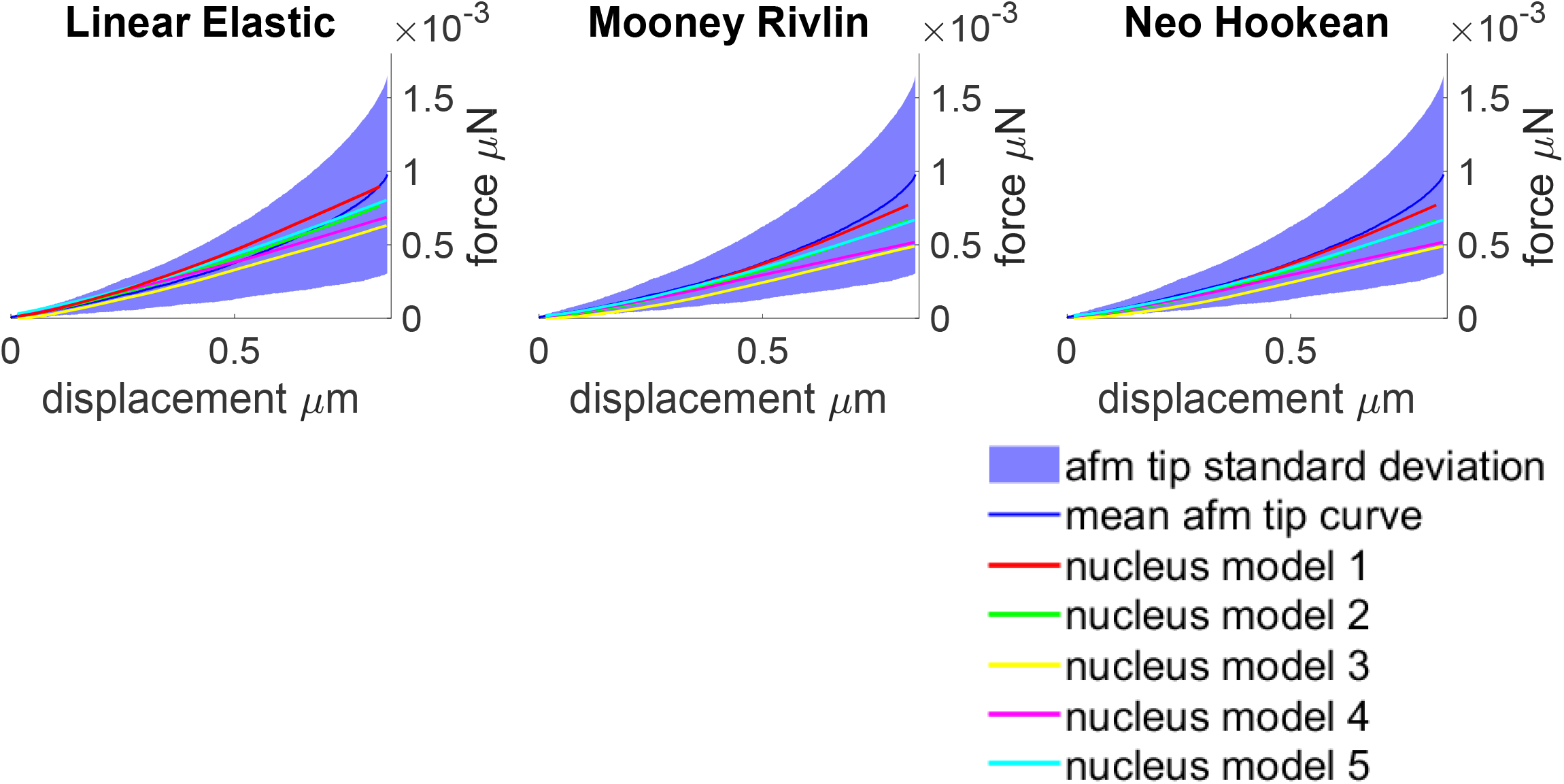
material elasticity comparison. Nuclei simulations of nucleus 4 and 5 were attempted using linear isotropic model that used the optimized values for the homogeneous image conversion(left). Mooney-Rivlin models were then created by converting the homogeneous optimized elasticity to Mooney-Rivlin constants as indicated within previous research^15^ by where c01 was set to 0 and c10 was formed by dividing the linear isotropic elasticity by 6(middle). From this point, 5 nuclei were formed by creating a Neo-Hookean material model by dividing the homogeneous optimized elasticity by 6 to form the Neo Hookean material constants as explained within the Abaqus user manual.

**Figure S3.**
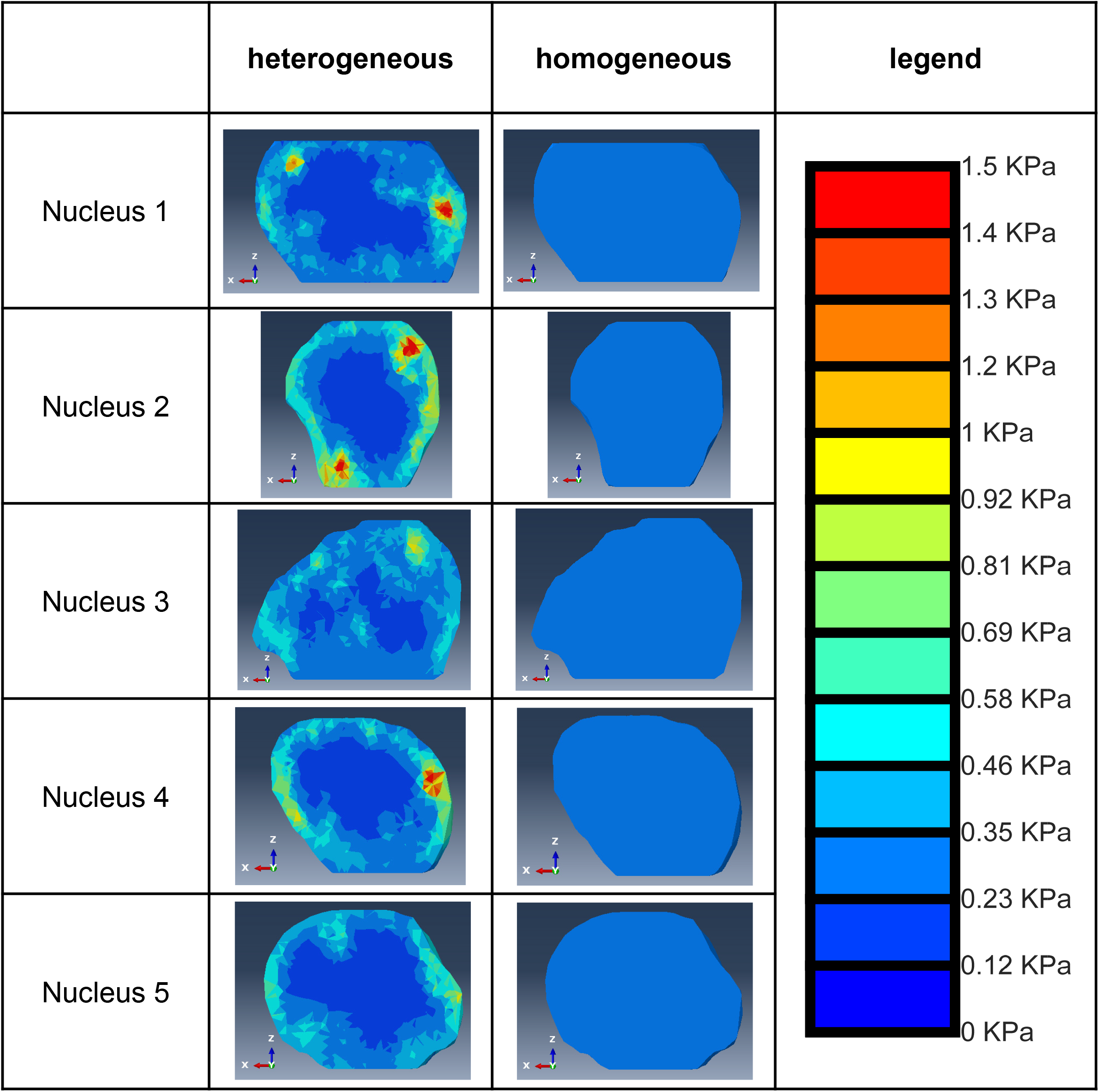
Chromatin material models. Finite element models of the nucleus have been defined with the chromatin images using both the homogeneous conversion factors and the heterogeneous conversion factors with a cross section of the nucleus models defined with the heterogeneous model shown on the left and cross sections of the homogeneous models shown on the right showing the material values using a color scale shown on the far right.

**Figure S4.**
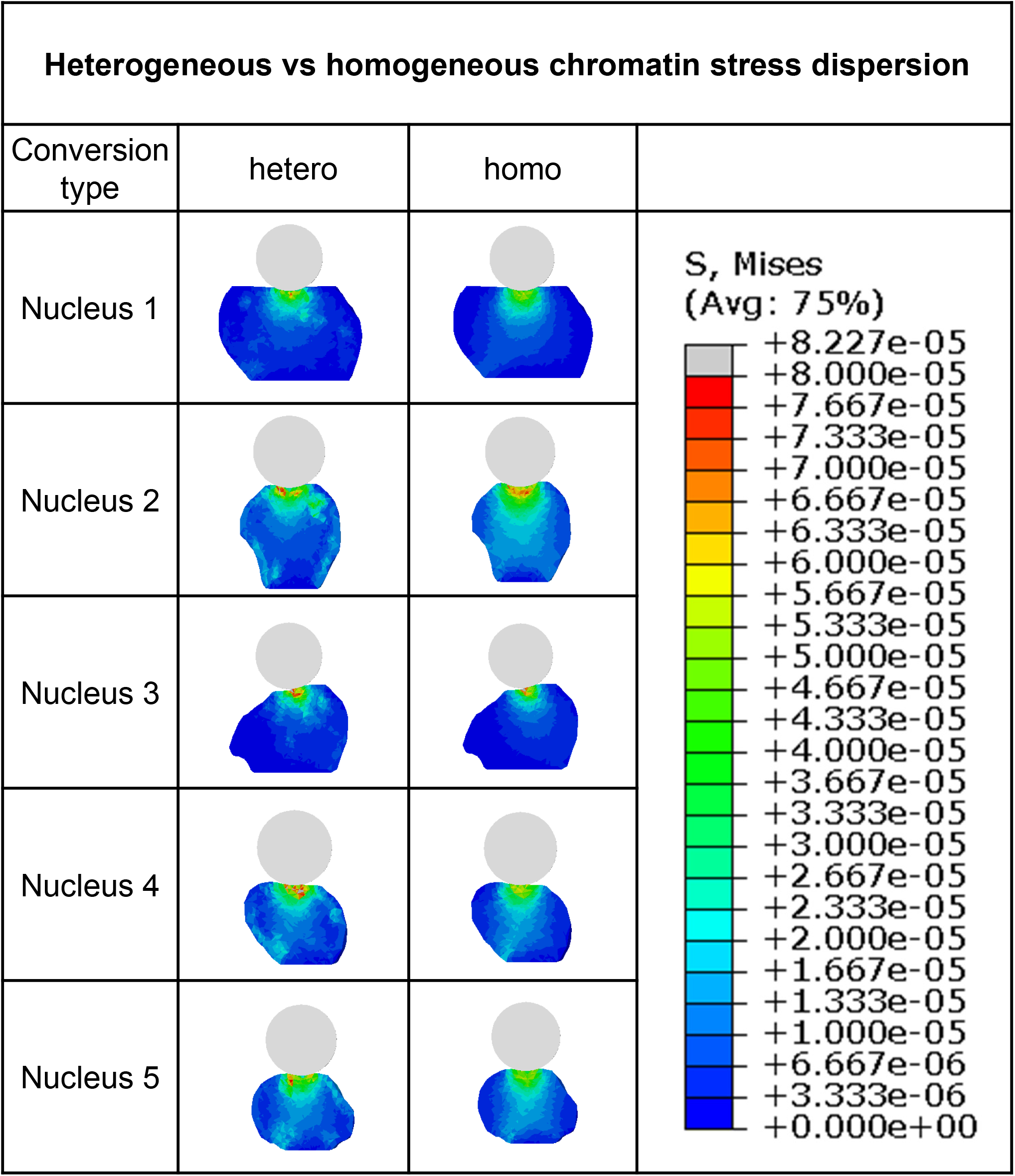
Heterogeneous vs Homogeneous stress distribution. Nuclei models 1-5 were created with optimized heterogeneous and homogeneous conversion factors. The nucleus models then underwent a simulated atomic force microscopy experiment with nucleus cross sections for the heterogeneous nuclei showing stress dispersions dependent on the chromatin density within the original images (left) as well as homogeneous models showing a stress dispersion not dependent on the original chromatin density (right).

**Figure S5.**
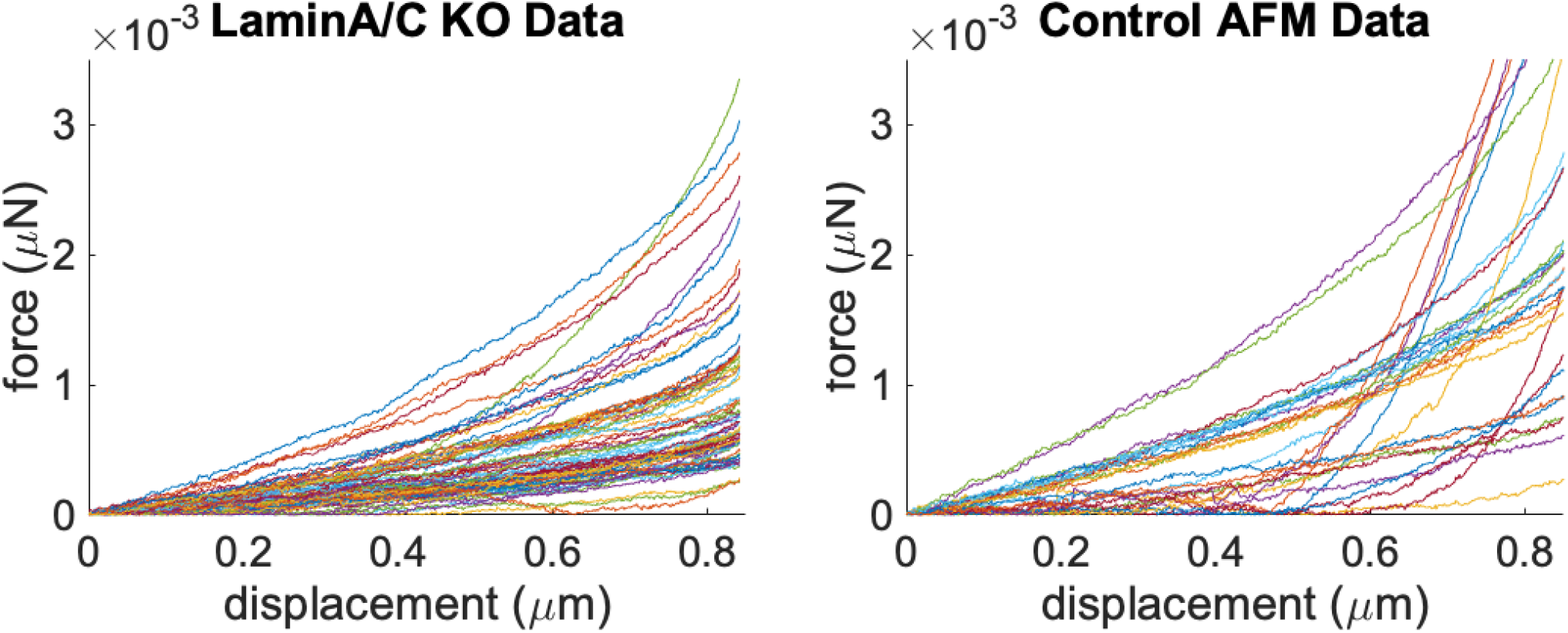
Atomic Force Microscopy Curves. Atomic force microscopy experiment data for LaminA/C knockdown nuclei (n=72, left) and control nuclei (n=30, right) has been plotted using Matlab.

